# On the role of cell chaining in the attenuation of a *Listeria monocytogenes divIVA* mutant

**DOI:** 10.64898/2026.01.05.697648

**Authors:** Sabrina Wamp, Gudrun Holland, Jeanine Rismondo, Sven Halbedel

## Abstract

*Listeria monocytogenes* is a facultative intracellular human pathogen capable of invading non-phagocytic host cells, replicating within their cytosol, and spreading directly from cell to cell. These processes are mediated by specialized virulence factors but also depend on the DivIVA protein. DivIVA aids in the secretion of peptidoglycan-degrading autolysins in a process that requires the accessory secretion ATPase SecA2, thereby promoting daughter cell separation following cytokinesis. Consequently, a Δ*divIVA* mutant forms elongated chains of unseparated daughter cells, which may explain its attenuated virulence. To further explore the role of cell chaining for attenuation, we here investigated how different morphological aberrations affect the virulence of *L. monocytogenes*. To this end, we generated coccoid *mreB* and filamentous *ezrA* mutants, and compared them to the Δ*divIVA* mutant in different *in vitro* infection assays. Coccoid or filamentous morphologies did not impair host cell invasion or intracellular replication, unlike cell chaining of the Δ*divIVA* mutant. Introduction of a hyperactive allele of the PrfA virulence regulator, resulting in constitutive overexpression of virulence genes, was sufficient to restore the invasion defect of the Δ*divIVA* mutant, despite its pronounced cell chaining phenotype, but did not recover intracellular replication. We isolated suppressors of the Δ*divIVA* mutant carrying mutations in *secA2*, which likely enhance the SecA2 ATPase activity. In these suppressors, autolysin secretion, daughter cell separation, and invasion were fully restored, and intracellular replication was partially recovered. Thus, maintaining normal rod-shaped morphology plays a minor role in *L. monocytogenes* pathogenesis. Instead, virulence attenuation in the Δ*divIVA* mutant is better explained by distortions in PrfA- and SecA2-dependent processes.

**Author Summary:** *Listeria monocytogenes* infects humans by entering host cells, replicating intracellularly, and spreading from infected cells to neighboring cells. The bacterium is rod-shaped and normally occurs as single or double cells that can efficiently complete all stages of infection. However, a class of virulence-attenuated *L. monocytogenes* mutants forms long cell chains, suggesting a link between cell morphology and virulence. One such mutant lacks the cell division protein DivIVA and is severely impaired in infection, as it neither invades host cells, replicates in the cytoplasm nor spreads from cell to cell. To determine whether virulence attenuation is caused by cell chaining itself or by additional functional defects, we compared the virulence of the *divIVA* mutant with that of mutants exhibiting other morphological abnormalities. In addition, we isolated suppressor mutations that restore normal cell separation in the *divIVA* mutant and examined their effects on virulence. Our results show that cell morphology per se does not generally determine listerial virulence. Instead, they indicate that specific virulence-related functions are impaired in chain-forming *L. monocytogenes* mutants. These findings advance our understanding of the virulence defects associated with cell chaining and provide a basis for identifying their real underlying molecular causes.

## Introduction

*Listeria monocytogenes* is a rod-shaped Gram-positive, foodborne pathogen capable of causing severe infections in elderly and immunocompromised individuals, with fatality rates reaching up to 30% [1, 2]. It shifts from a saprophytic lifestyle in the environment to a pathogenic state upon entering the host, activating virulence factors while adapting key processes like metabolism and cell division [3]. The central transcription factor, PrfA, activates virulence gene expression during infection [4, 5]. The PrfA regulon includes *inlAB* genes, which encode internalins A and B essential for host cell invasion [6, 7], and several virulence genes within the LIPI-1 pathogenicity island [8]. These include *hly*, *plcA*, and *plcB*, which encode the hemolysin listeriolysin O and two phospholipases that disrupt phagosomal membranes [9–11], as well as *actA*, which encodes a surface protein crucial for cell-to-cell spread [12, 13]. After escaping the phagosome, ActA recruits host actin to the bacterial surface, forming tail-like structures that allow intracellular and intercellular movement and promote infection of neighboring cells [14, 15]. By coating itself with host actin, the bacteria can also evade autophagic recognition [16].

For efficient invasion, intracellular replication, and cell-to-cell spread, *L. monocytogenes* also depends on house-keeping proteins, such as DivIVA [17, 18]. DivIVA is a late cell division protein found in many Gram-positive bacteria, where it binds to curved membranes at the cell poles and the invaginating membrane regions of the division septum [19–21]. DivIVA coordinates various cellular processes across different species, including cell division, sporulation, swarming motility, and biofilm formation, typically by recruiting protein-binding partners to the poles and the septum [22, 23]. Cells of a *L. monocytogenes* Δ*divIVA* mutant fail to separate after cell division leading to the formation of long cell chains that are unable to invade, replicate intracellularly or spread from cell to cell [17, 24]. Cell chaining of the Δ*divIVA* mutant is explained by the lack of secretion of the two autolysins, CwhA (p60) and NamA [17], which hydrolyze peptidoglycan from the outside of the division septum [25–27]. Secretion of CwhA and NamA requires the accessory secretion ATPase SecA2 [27, 28]. The role of DivIVA in SecA2-dependent autolysin secretion is not fully understood, but it has been shown that the preproteins of CwhA and NamA are sequestered to the septa by DivIVA, where SecA2 is also located [17].

Daughter cell separation in chain-forming *L. monocytogenes* mutants can be restored by addition of exogenous autolysin sources or by mechanical disruption restoring some but not all virulence functions [17, 29–31]. This has shaped the notion that the virulence defect of chain-forming mutants is mainly due to morphological reasons. Changes in cell shape are known to influence infection-related processes in other species. In uropathogenic *Escherichia coli*, filamentous forms evade phagosomal killing by macrophages as they are harder to engulf [32]. On the other hand, an elongated bacterial surface can facilitate host cell attachment, as reported for filamentous forms of the lung pathogen *Legionella pneumophila* [33]. Similarly, *Streptococcus pneumoniae* can grow in chains of different length with short chains enabling complement evasion and long chains improving host cell attachment [34].

To better understand the role of cell chaining in the attenuation of the Δ*divIVA* mutant, we wanted to analyze how different cell morphologies impact the virulence of *L. monocytogenes*. We therefore generated strains allowing depletion of EzrA, a negative regulator of FtsZ ring formation, and MreB, an actin-like cytoskeleton protein. EzrA links the FtsZ ring to the cytoplasmic membrane [35], while MreB provides a scaffold that generates the spatial arrangement and circumferential movement of peptidoglycan-synthesizing elongasomes along the long axis of rod-shaped bacteria [36, 37]. The *ezrA* and *mreB* genes are essential in *L. monocytogenes* [38, 39], and EzrA depletion resulted in the formation of elongated *L. monocytogenes* cells [38]. MreB is the only essential protein out of three actin-like paralogs of *L. monocytogenes* [39]. In rod-shaped *B. subtilis*, MreB removal leads to cell shortening with an inflated, rounded morphology [40] or even complete collapse of rod-shape, resulting in spherical cells [41]. We here described the morphological consequences of EzrA and MreB depletion in *L. monocytogenes* and compare the virulence of these depletion strains to the Δ*divIVA* mutant in various *in vitro* infection assays. Mutations in other genes selectively suppressing the different virulence defects of the Δ*divIVA* mutant are presented.

## Results

### Distinct morphological phenotypes of *L. monocytogenes divIVA*, *ezrA* and *mreB* mutants

For investigation of the role of cell morphology on *L. monocytogenes* virulence, we generated mutants with different morphological defects. For this, we first attempted to delete the *mreB* gene (*lmo1548*), but without success. However, the *mreB* gene could be deleted on IPTG-containing plates in a strain that carried an IPTG-inducible copy of *mreB* at an ectopic site. The resulting MreB depletion strain (LMSW49) (genotype designated as i*mreB*), grew as the wild type when the inoculum was taken from a preculture grown in the presence of IPTG, but only grew slowly when the preculture was grown without the inducer (compare 1^st^ and 2^nd^ depletion, Fig. 1A). It is important to note here that MreB levels were already reduced during the first round of depletion as evidenced by Western blotting (Fig. 1B), even though no growth defect became apparent at this stage of depletion (Fig. 1A). Strain LMSW49 rapidly accumulated suppressors on BHI agar plates not containing IPTG, but this growth defect was neutralized when 0.3 M sucrose was added as an osmo-protectant, allowing strain LMSW49 to grow on BHI agar even in the absence of IPTG (data not shown). When sucrose was added to the agar, i*mreB* cells formed rods in the presence of IPTG but entirely lost their rod shape and formed coccoid cells instead when no IPTG was present (Fig. 1C).

**Figure 1:**
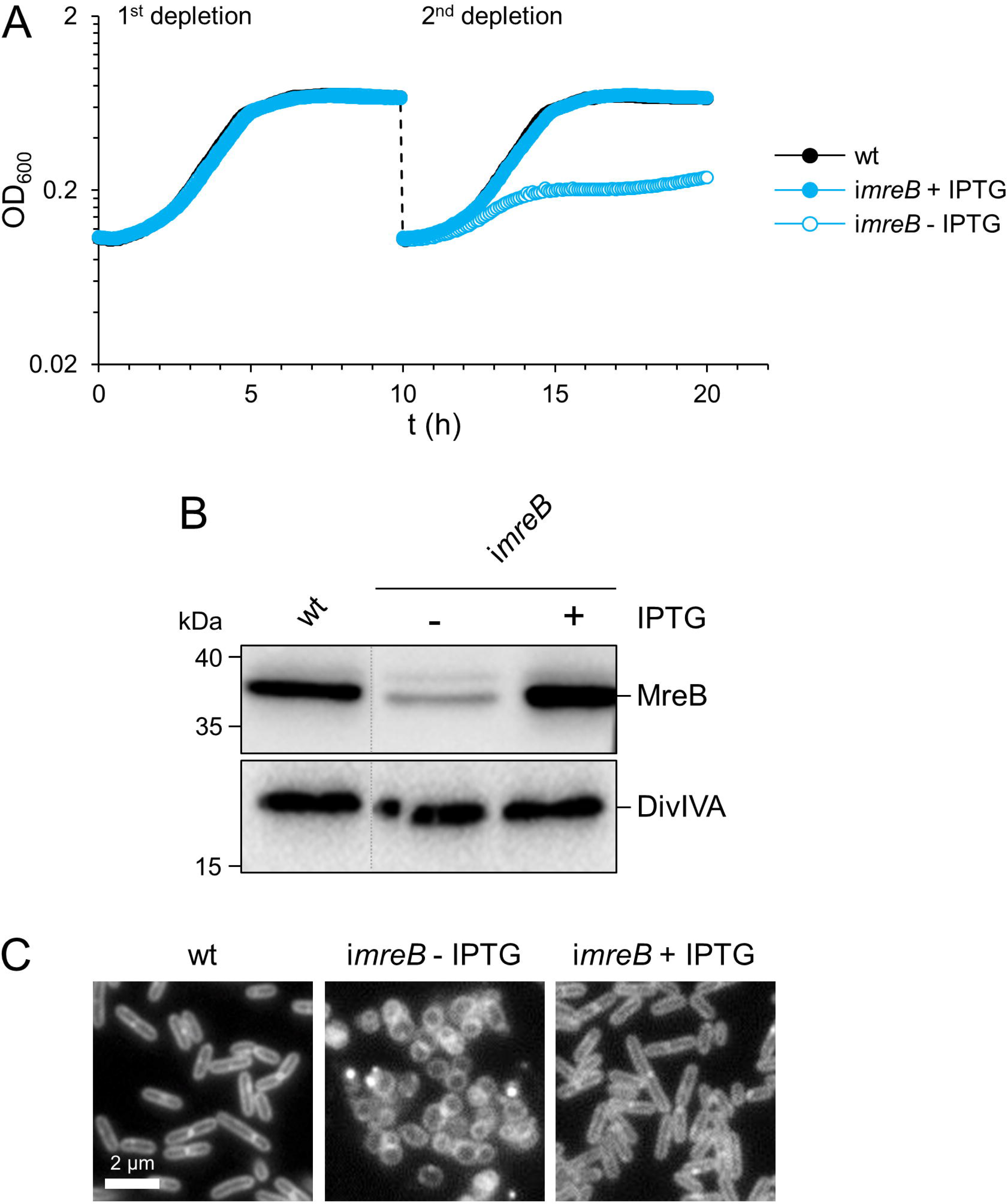
Depletion of MreB in *L. monocytogenes*. (A) Effect of MreB depletion on growth of *L. monocytogenes*. Strains EGD-e (wt) and i*mreB* (LMSW49) were grown in BHI broth ± 1 mM IPTG. Although this did not immediately result in a growth defect, a growth defect became apparent in the second depletion when new cultures were started with the MreB-depleted stationary phase cells from the first depletion as inoculum. The experiments were independently repeated three times and one representative set of experiments is shown. (B) Western blot showing MreB (upper panel) and DivIVA (lower panel) levels in *L. monocytogenes* strains EGD-e (wt) and LMSW49 (i*mreB*) grown in BHI broth ± 1 mM IPTG at 37°C in mid-exponential growth phase during a cultivation similar to the first depletion shown in panel A. Dotted lines mark the positions at which irrelevant lanes were removed from the blots. (C) Micrographs showing the cellular morphology of the same set of strains during growth on BHI agar plates containing 0.3 M sucrose. 1 mM IPTG was added where indicated. Membranes were stained with nile red.

Next, we tried to generate a filamentous mutant and attempted deletion of *ezrA*, required for cell division [38], but this was also not possible in wild type background. However, as shown in a previous report [38], *ezrA* could be deleted when a second IPTG-dependent *ezrA* copy was present. Similar to the i*mreB* depletion strain, the i*ezrA* (LMJR183) mutant did not require IPTG for growth when the inoculum was taken from a preculture grown in the presence of IPTG; however, it grew in an IPTG-dependent manner, when a depleted preculture was used as the inoculum (Fig. 2A).

**Figure 2:**
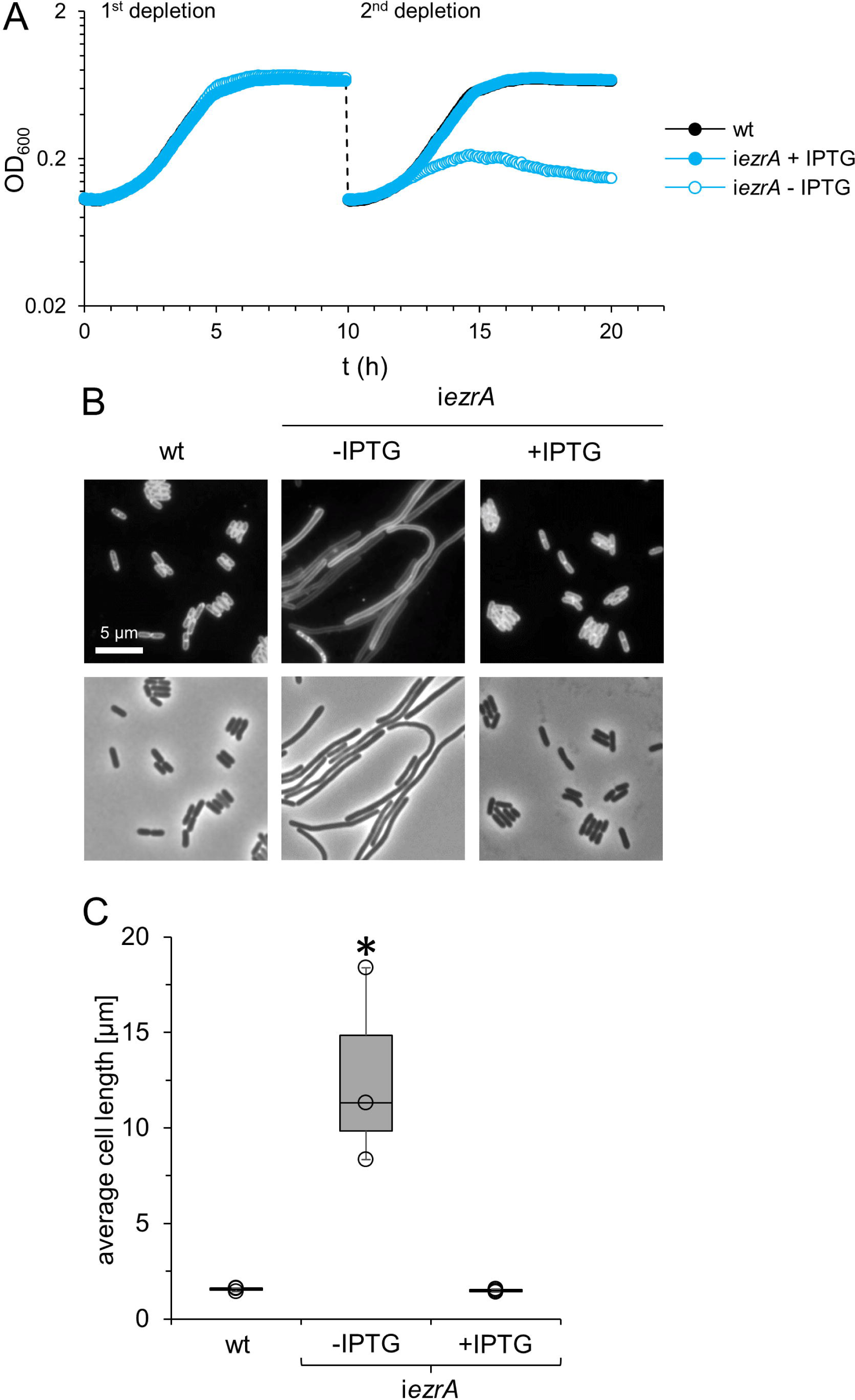
EzrA depletion affects growth and cell division of *L. monocytogenes*. (A) Effect on growth. Strains EGD-e (wt) and LMJR183 (i*ezrA*) were grown in BHI broth ± 1 mM IPTG at 37°C. The i*ezrA* cultures were started at 0 h in fresh BHI broth ± IPTG using a preculture grown in the presence of 1 mM IPTG. Although this did not immediately result in a growth defect, a growth defect became apparent in the second depletion when new cultures were started with the EzrA-depleted stationary phase cells from the first depletion as inoculum. The experiments were independently repeated three times and one representative set of experiments is shown. (B-C) Effect on cell length. Micrographs showing cells taken from the first depletion as in panel A after nile red staining (B) and a box blot showing their average cell lengths (C). Average cell lengths from three experiments with 300 measured cells per experiments are shown. Statistical significance is indicated (* - *P*=0.02, *t*-test).

Importantly, the expected *ezrA* phenotype was already evident during the first round of depletion as severe filamentation was observed in the absence of IPTG (Fig. 2B-C). EzrA-depleted cells were on average 12.7±5.2 µm long (wild type cell length: 1.5±0.1 µm, Fig. 2B-C), and could form filaments as long as 30-50 µm without a single septum. These results are is in agreement with a previous Tn-Seq study demonstrating essentiality of *mreB* and *ezrA* [39]. In contrast to the *mreB* and *ezrA* phenotypes, a mutant lacking the *divIVA* cell division gene grew as fast as the wild type (Fig. 3A), but formed long chains of cells (Fig. 3B-C) consistent with previous reports [17, 18].

**Figure 3:**
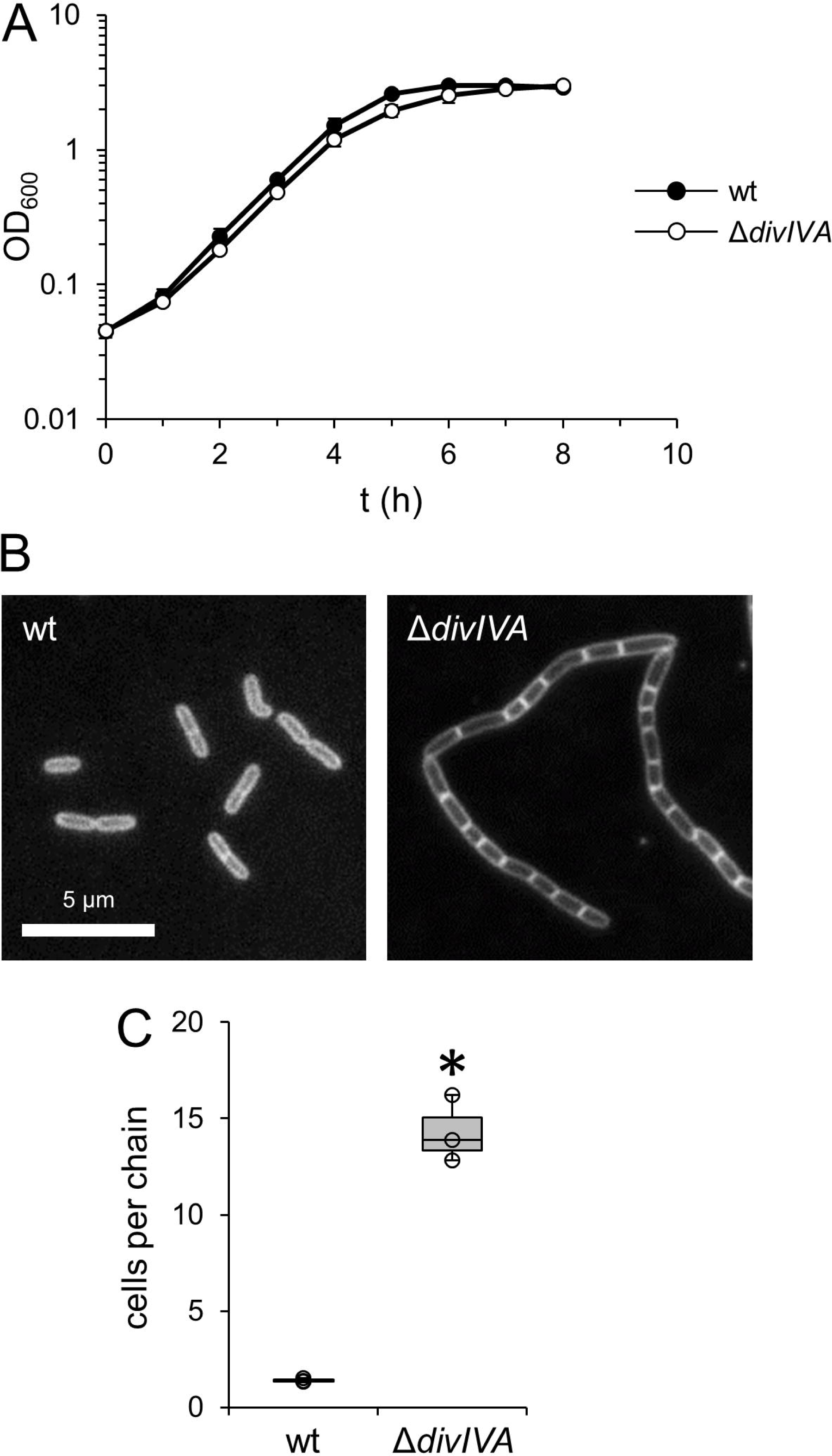
Effect of *divIVA* deletion on growth and cell division of *L. monocytogenes*. (A) Growth of *L. monocytogenes* strains EGD-e (wt) and LMS2 (Δ*divIVA*) in BHI broth at 37°C. Average values and standard deviations were calculated from three independent repetitions. (B) Micrographs showing morphology of the same strains during mid-logarithmic growth after nile red staining. (C) Number of cells per chain. Average values from three experiments with 40 chains (Δ*divIVA*) or singlet/doublets (wt) each are shown. The asterisk indicates statistical significance (*P*<0.01, *t*-test).

Scanning electron microscopy further confirmed characteristic morphological differences between *divIVA*, *ezrA* and *mreB* mutants. The Δ*divIVA* mutant formed filamentous cells with occasional gentle circular wall constrictions likely representing putative division sites. Some of these filamentous cells were locally bent or even coiled. The surface structure was similar as in wild type but regularly showed small vesicle-like structures (Fig. 4, Fig. S1). Coiling and small vesicle-like structures at the wall surface were generally not observed in EzrA-depleted cells (Fig. 4, Fig. S1). In contrast, cells depleted for MreB were spherical and revealed two clearly distinct zones of surface ultrastructure: (1) a smooth zone likely representing septal peptidoglycan and (2) a rough zone with granular to particulate elevations of peptidoglycan (Fig. 4, Fig. S1).

**Figure 4:**
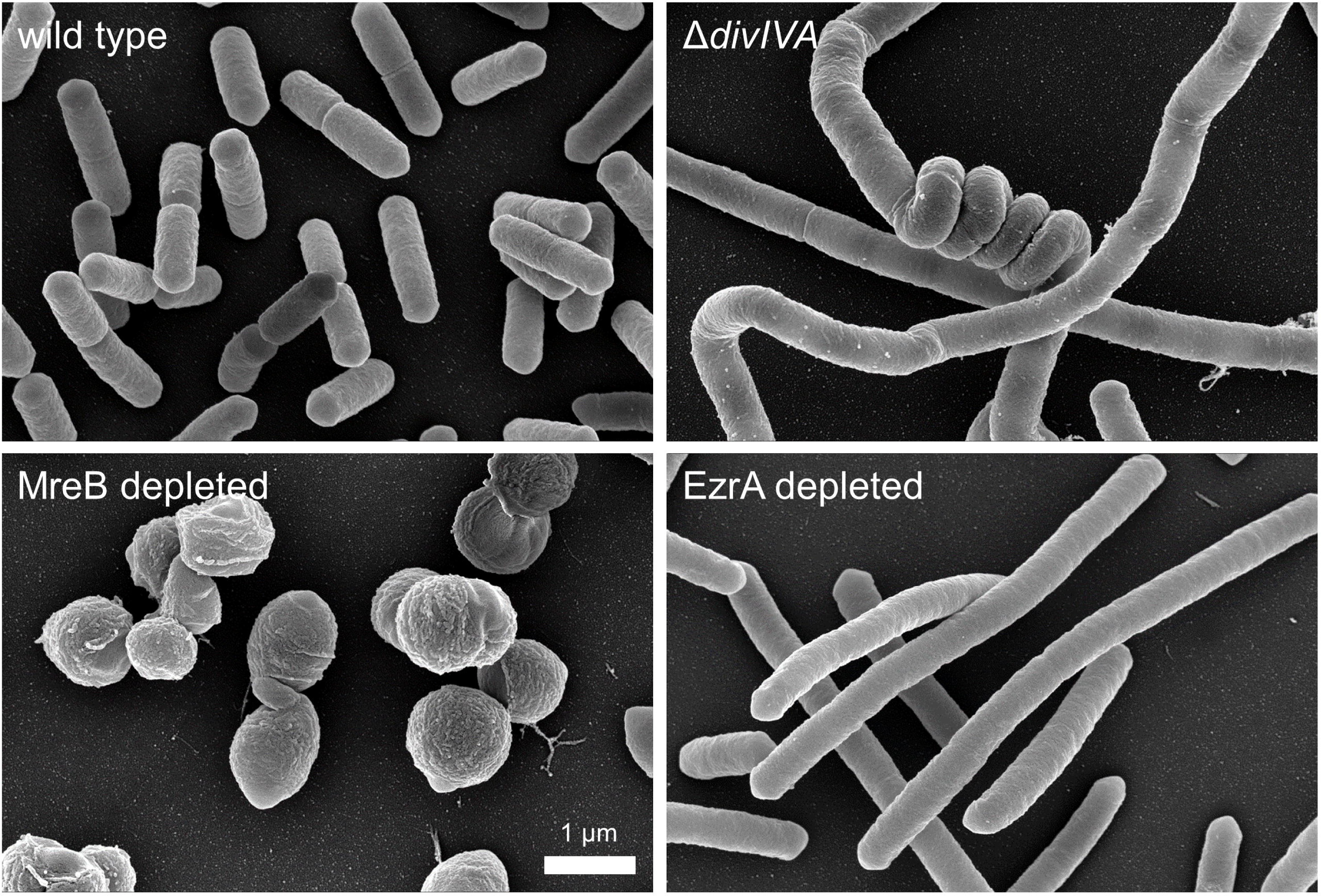
Ultrastructure of *L. monocytogenes divIVA*, *ezrA* and *mreB* mutants. Scanning electron microscopy images of *L. monocytogenes* strains EGD-e (wild type), LMS2 (Δ*divIVA*), LMJR183 (i*ezrA*) and LMSW49 (i*mreB*). Strains were cultivated in BHI broth at 37°C to mid-logarithmic growth. For the IPTG-dependent strains, the preculture contained IPTG, but the inducer was omitted in the main culture.

### Combination of *divIVA*, *ezrA* and *mreB* mutations

We wondered whether the combination of *divIVA*, *ezrA* and *mreB* mutations would generate additional morphotypes and deleted *divIVA* in i*ezrA* and i*mreB* backgrounds. In contrast to the growth behavior of the parental i*ezrA* and i*mreB* strains (Fig. 1A, Fig. 2A), IPTG-dependent growth of the resulting strains became already evident during the first round of EzrA- or MreB-depletion (Fig. S2), indicating that the *divIVA* deletion somehow exacerbates *ezrA* and *mreB* phenotypes. For inspection of the morphology of *divIVA mreB* cells, MreB was depleted from strain LMSW207 (Δ*divIVA* i*mreB*). Fluorescence microscopy of membrane stained cells revealed mainly streptococcoid morphology under this condition with formation of chains of cocci (Fig. 5A). Scanning electron microscopy showed that the individual cells of these chains were of variable size, occasionally formed protrusions and depositions and frequently underwent lysis, as evidenced by holes in their sacculi. (Fig. 5B). In contrast, depletion of EzrA from LMSW208 cells (Δ*divIVA* i*ezrA*) yielded chains of filaments, which was expected (Fig. 5A). As described for the Δ*divIVA* single mutant above, these filaments also generated coiled regions (Fig. 5A-B). EzrA-depleted Δ*divIVA* cells also regularly showed signs of lysis (*i. e.* flaccid or swollen cells). We did not consider these mutants for further analyses due to their lytic phenotypes.

**Figure 5:**
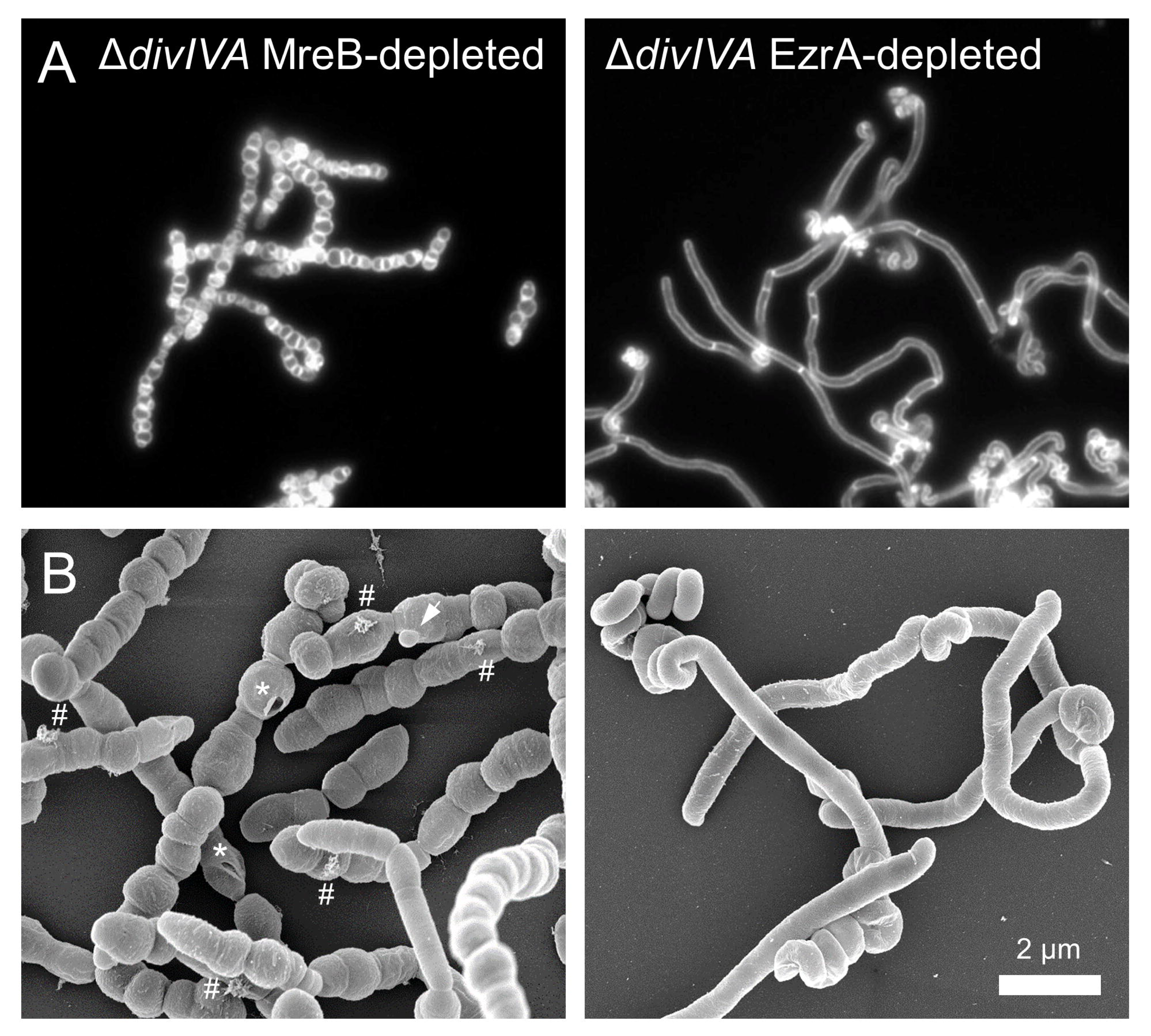
Combination of *ezrA*, *divIVA* and *mreB* phenotypes. (A) Micrographs showing nile red stained cells of strains LMSW207 (Δ*divIVA* i*mreB*) and LMSW208 (Δ*divIVA* i*ezrA*) after cultivation in BHI broth not containing IPTG. (B) Scanning electron microscopy images of the same strains. Distortions in the cell wall ultra-structure are indicated by asterisks (holes), arrows (protrusions) and hash marks (depositions).

### Effects of morphological aberrations on host cell invasion, intracellular growth and cell-to-cell spread

To determine the effects of chain forming, coccoid and filamentous growth on the ability of *L. monocytogenes* to invade non-phagocytic cells, human HepG2 hepatocytes were infected with the Δ*divIVA* mutant as well as with MreB and EzrA depletion strains. Invasion of hepatocytes is clearly PrfA-dependent (Fig. 6A) reflecting the known involvement of PrfA-dependent virulence genes in hepatocyte invasion [7, 42]. For the Δ*divIVA* mutant, we observed a 15-fold reduction of invasion rate, which also is in good agreement with a previous report [17]. However, no such reduction of invasion rate was observed with cells depleted for MreB or EzrA (Fig. 6A), which supports the idea that the aberrant morphology of the Δ*divIVA* mutant likely does not suffice to explain its invasion defect.

**Figure 6:**
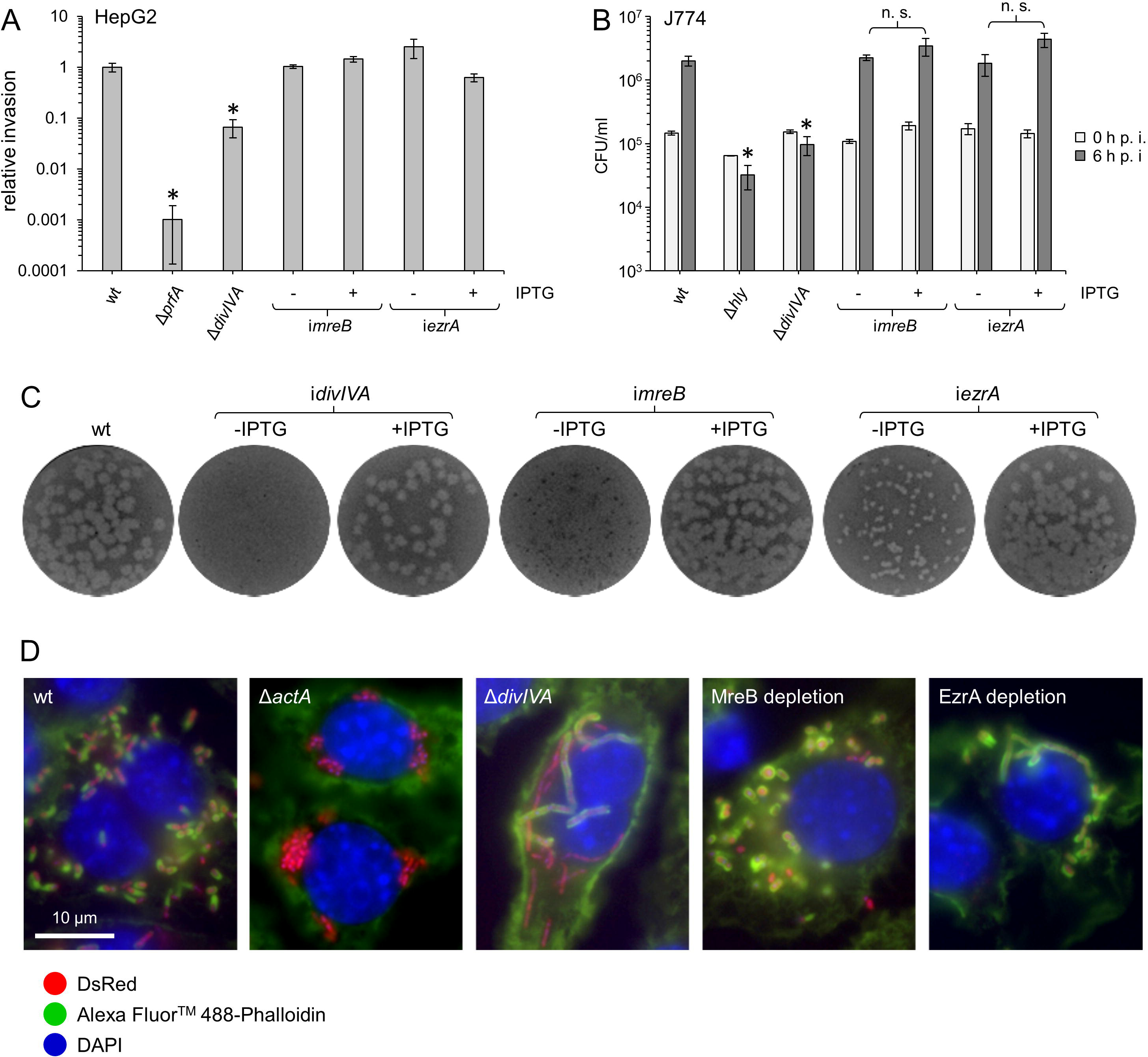
Effect of morphological aberrations on invasion, intracellular replication, cell-to-cell spread and actin tail formation. (A) Contribution of *divIVA*, *mreB* and *ezrA* to hepatocyte invasion. Invasion of *L. monocytogenes* strains EGD-e (wt), BUG2214 (Δ*prfA*), LMS2 (Δ*divIVA*), LMSW49 (i*mreB*) and LMJR183 (i*ezrA*) pre-grown in the absence or presence of 1 mM IPTG into HepG2 hepatocytes. Values are expressed relative to wild type. Average values and standard deviations are shown. Asterisks mark statistically significant differences compared to wild type (*P*<0.01, *t*-test with Bonferroni Holm correction). (B) Contribution of *divIVA*, *mreB* and *ezrA* to intracellular growth in macrophages. J774 mouse macrophages were infected with *L. monocytogenes* strains EGD-e (wt), LMS250 (Δ*hly*), LMS2 (Δ*divIVA*), LMSW49 (i*mreB*) and LMJR183 (i*ezrA*) and then cultivated in the absence or presence of IPTG, where appropriate. Average values and standard deviations were calculated from experiments performed in triplicates. Statistically significant differences (compared to wild type or between depleted and induced conditions) are marked by asterisks (*P*<0.01, *t*-test with Bonferroni-Holm correction, n. s. – not significant). (C) Effect of DivIVA, MreB or EzrA depletion on cell-to-cell spread. Plaque formation assay using 3T3 mouse fibroblasts with *L. monocyctogenes* strains EGD-e (wt), LMS30 (i*divIVA*), LMSW49 (i*mreB*) and LMJR183 (i*ezrA*) ± IPTG. (D) Necessity of DivIVA, MreB and EzrA for actin coating and actin tail formation. J774 mouse macrophages were infected with DsRed producing *L. monocytogenes* strains LMJD20 (wt), LMSF2 (Δ*actA*), LMSW205 (Δ*divIVA*), LMSW202 (EzrA depletion) and LMSW203 (MreB depletion). Infections were carried out in the absence of IPTG and analyzed by fluorescence microscopy 4 hours post infection after DAPI and phalloidin staining. Composite images are shown.

We next determined the role of DivIVA, EzrA and MreB in intracellular growth in mouse macrophages. To this end, IPTG-dependent i*ezrA* and i*mreB* strains were pre-grown with IPTG and then used to infect J774 mouse macrophages, which were then further incubated either in the presence or absence of IPTG. Surprisingly, depletion of EzrA or MreB did not result in a significant reduction of the final intracellular bacterial loads in the infected macrophages six hours post infection. In stark contrast, deletion of *divIVA* prevented intracellular growth in mouse macrophages (Fig. 6B). In order to make sure that the expected morphological phenotypes were fully established intracellularly despite the short depletion period, we repeated the infection assay with strains carrying a plasmid encoding the red fluorescent marker DsRedExpress for microscopic analysis of bacterial morphology during infection. Infection with wild-type cells led to the detection of fluorescent rods in the cytoplasm of infected macrophages, whereas the Δ*divIVA* mutant formed chains, and fluorescent filaments and cocci were observed in the macrophage cytoplasm during intracellular EzrA and MreB depletion, respectively (Fig. S3). This showed that the expected morphological phenotypes were clearly established during infection and confirmed that growth with coccoid or filamentous morphology does not impair intracellular growth of *L. monocytogenes* inside macrophages.

To further study the effect of cell shape changes on virulence, we then determined the importance of DivIVA, EzrA and MreB for cell-to-cell spread in a plaque assay using 3T3 mouse fibroblasts. IPTG-dependent mutants grown with the inducer were used to ensure normal invasion, but IPTG was omitted after cell entry. As can be seen in Fig. 6C, depletion of DivIVA and MreB completely prevented plaque formation, whereas depletion of EzrA resulted in smaller plaques. Actin tail assembly is a main prerequisite for plaque formation [12, 13]. Therefore, J774 macrophages were infected with i*ezrA*, Δ*divIVA* and i*mreB* mutants expressing DsRedExpress and host actin was stained with fluorescent phalloidin 4 h post infection. Coating with actin and the formation of actin tails were readily observed in cells infected with the wild type, whereas neither actin coating nor tail formation was detected in cells infected with the Δ*actA* mutant, as expected. In cells depleted of MreB and EzrA, actin coating was present, but tail formation was only rarely observed. In contrast, infections with the Δ*divIVA* mutant revealed both actin-coated and uncoated cell chains and the complete absence of actin tails (Fig. 6D). These findings suggest that the Δ*divIVA* mutant is generally capable of reaching the host cytoplasm, but that phagosomal escape may be delayed or actin recruitment distorted.

### Suppression of the Δ*divIVA* invasion defect by PrfA activation

Since there is no clear correlation between morphology and virulence, other explanations are needed for the versatile virulence defects of the Δ*divIVA* mutant. We therefore concluded that DivIVA could be required for full activity or subcellular targeting of some of the classical virulence factors, the transcription of which is controlled by the transcription factor PrfA. To test this idea, we made use of the *prfA G145S* allele (*prfA\**), which encodes a PrfA variant that is constitutively active causing constitutive overexpression of PrfA-dependent virulence genes [43], to generate a Δ*divIVA* mutant with PrfA switched on. The resulting Δ*divIVA prfA\** strain has increased hemolytic and phospholipolytic activity just as the *prfA*\* strain, but is non-motile and forms cell chains just as the Δ*divIVA* mutant (Fig. S4). As invasion of HepG2 hepatocytes requires internalins A and B [7, 44], whose transcription is PrfA-dependent [45], invasion of a Δ*prfA* mutant into hepatocytes is reduced and that of a *prfA*\* mutant increased (Fig. 7A), due to variations in *inlAB* expression. Surprisingly, the invasion defect of the Δ*divIVA* mutant was not only abolished by the introduction of the *prfA\** mutation, but instead the Δ*divIVA prfA\** double mutant even exhibited the increased invasiveness of the *prfA\** single mutant (Fig. 7A). In stark contrast to this, introduction of the *prfA\** mutation into the Δ*divIVA* mutant did not cause suppression of the intracellular replication defect of the Δ*divIVA* mutant in J774 macrophages (Fig. 7B) or its spreading defect in mouse fibroblasts (Fig. 7C). This unequivocally demonstrates that (i) hepatocyte invasion is generally not restricted by chain formation, (ii) the invasion defect of the Δ*divIVA* mutant involves a PrfA-dependent process, and (iii) the intracellular replication and cell-to-cell spread defects of the *divIVA* mutant cannot be explained by malfunction of a PrfA-dependent virulence factor.

**Figure 7:**
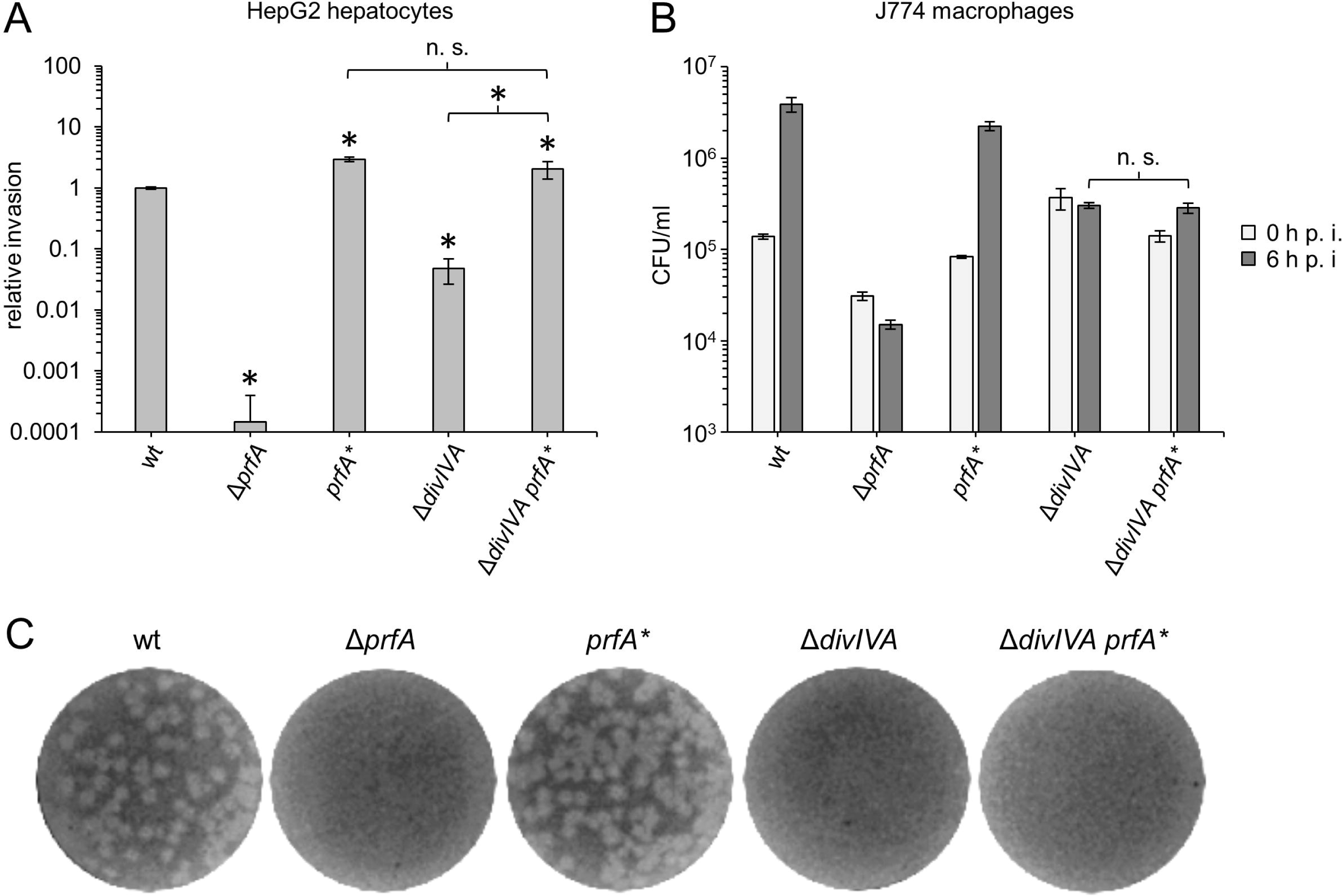
Differential effects of the *prfA*\* mutation on invasion and intracellular multiplication of the Δ*divIVA* mutant. (A) The *prfA\** mutation suppresses the invasion defect of the Δ*divIVA* mutant. Invasion of *L. monocytogenes* strains EGD-e (wt), BUG2214 (Δ*prfA*), BUG3057 (*prfA\**), LMS2 (Δ*divIVA*) and LMSW251 (Δ*divIVA prfA\**) into HepG2 hepatocytes. Values are expressed relative to wild type. Average values and standard deviations are shown. The asterisks mark statistically significant differences compared to wild type (*P*<0.01, *t*-test with Bonferroni-Holm correction) or to the Δ*divIVA prfA\** mutant (*P*<0.01, *t*-test, n. s. – not significant). (B) No suppression of the intracellular replication defect of the Δ*divIVA* mutant by the *prfA\** mutation. J774 mouse macrophages were infected with the same strains as in panel A. Average values and standard deviations were calculated from experiments performed in triplicates. No statistically significant difference in the number of intracellular bacteria was detected between Δ*divIVA* and Δ*divIVA prfA\** cells six hours post infection (*P*<0.01, *t*-test, n. s. – not significant). (C) Dominance of the Δ*divIVA* deletion over the *prfA\** mutation in a plaque forming assay. The same strains as above were used to infect 3T3 mouse fibroblasts to analyze cell-to-cell spread.

### Suppression of the Δ*divIVA* intracellular replication defect by SecA2 activation

In rare cases, we noticed the occurrence of smooth suppressors in isolation streaks of the Δ*divIVA* mutant, which forms rough colonies on BHI agar plates. We isolated two such spontaneous smooth Δ*divIVA* suppressors, designated *ssd1* and *ssd2* (<u>s</u>mooth <u>s</u>uppressor of Δ*<u>d</u>ivIVA*). Whole genome sequencing demonstrated that *ssd1* had a M95I exchange in the *secA2* gene, encoding the accessory preprotein translocase [28], whereas *ssd2* carried three mutations in addition to the Δ*divIVA* deletion: A T123A exchange in *secA2,* a G98V substitution in *tagH*, encoding the ATP binding component of an ABC transporter for teichoic acid translocation, and a frameshift in *eslA*, coding for the ATP binding component of an ABC transporter linked to lysozyme resistance [46], that causes an I229Y substitution followed by a premature stop codon four codons downstream. Since the *secA2 M95I* substitution was the only genetic alteration found in the *ssd1* suppressor and given that DivIVA is required for SecA2-dependent protein secretion in *L. monocytogenes* [17], we concluded that the *secA2* mutations represent the relevant genetic adaptation responsible for suppressing the rough phenotype. In addition, the M95I substitution affects a conserved methionine in the Walker A motif, which is present in all SecA proteins and is essential for ATP-binding and hydrolysis [47]. The T123A mutation in suppressor *ssd2* also maps to the nucleotide binding domain of SecA2 (Fig. 8A) and is even identical to the classical *azi-1* mutation, conferring azide resistance to housekeeping *B. subtilis* SecA by activation of its ATPase activity [48]. Therefore, the *secA2 M95I* and *T123A* mutations were recreated in the Δ*divIVA* background by allelic exchange. Microscopic analysis showed that the Δ*divIVA secA2 M95I* and the Δ*divIVA secA2 T123A* mutants formed doublets and individual cells instead of long cell chains (Fig. 8B). Separation of exoproteomes consistently revealed that both strains secreted CwhA (p60, Fig. 8C), which is the most prominent band in *L. monocytogenes* exoproteomes. Despite suppression of cell chaining, motility on soft agar was not restored by either of the two *secA2* mutations (Fig. 8D), indicating that the function of DivIVA in flagellar motility is distinct from its role in SecA2-dependent protein secretion. However, when cell-to-cell spread was analyzed in a plaque assay, partial suppression of the plaquing defect of the Δ*divIVA* mutant was observed (Fig. 8E), indicating that invasion and/or intracellular replication defects of the Δ*divIVA* mutant can be compensated by specific *secA2* mutations. Separate assays of invasion and intracellular replication showed full suppression of the Δ*divIVA* mutant ’s defect in hepatocyte invasion (Fig. 8F) and partial suppression of its inability to replicate inside macrophages (Fig. 8G). These results demonstrate that individual virulence defects of the Δ*divIVA* mutant can be suppressed by specific amino acid substitutions in the nucleotide binding domain of SecA2.

**Figure 8:**
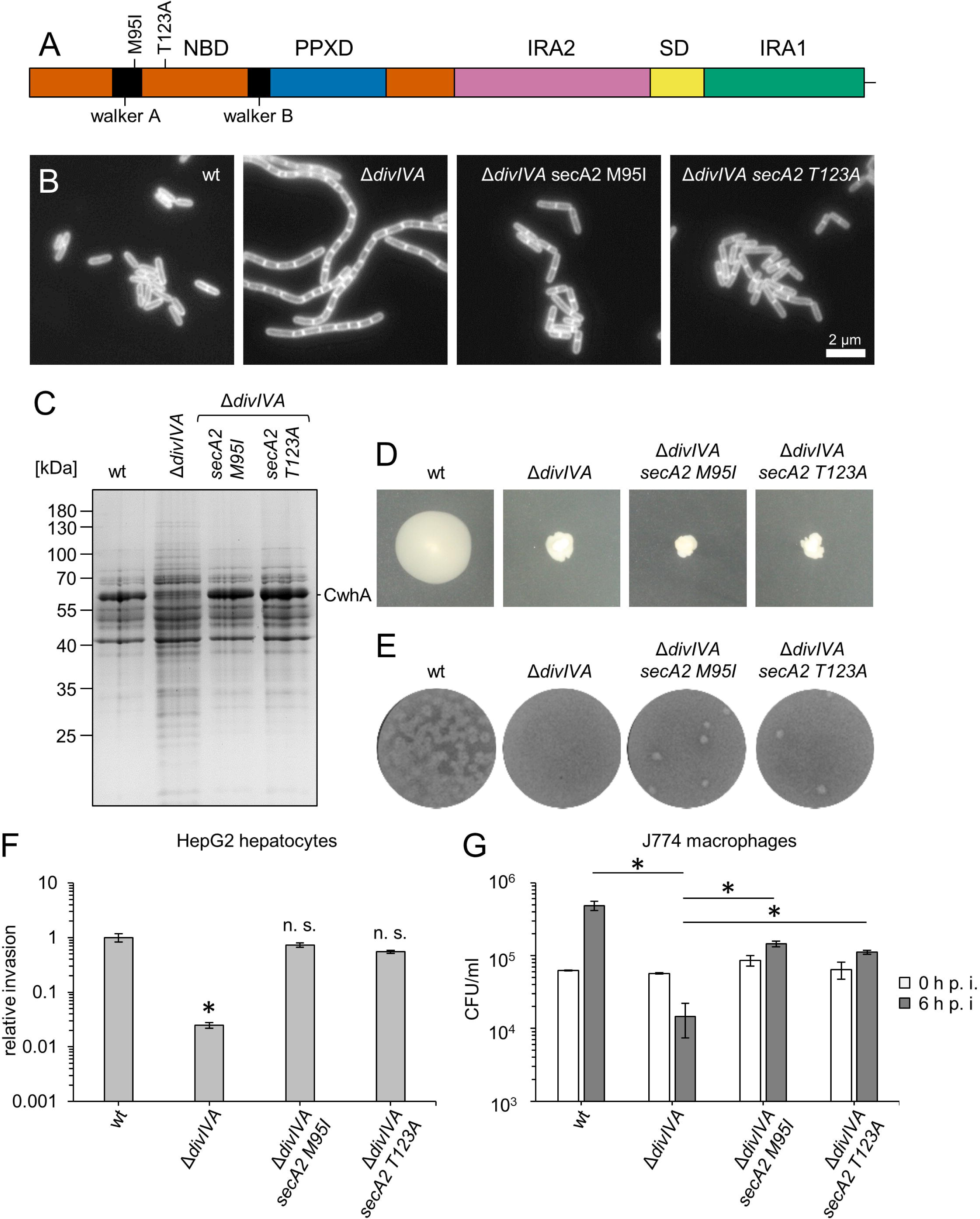
Partial virulence restoration in the Δ*divIVA* mutant by *secA2* mutations. (A) Localisation of Δ*divIVA* suppressor mutations M95I and T123A in the SecA2 protein. SecA2 domains (colored) and walker A and B motifs (black) are indicated. Abbreviations: NBD –nucleotide binding domain, PPXD – preprotein crosslinking domain, IRA 1/2 – intramolecular regulator of ATPase domains 1 and 2, SD – scaffold domain. (B) Micrographs showing the cellular morphology of *L. monocytogenes* strains EGD-e (wt), LMS2 (Δ*divIVA*), LMSW222 (Δ*divIVA secA2 M95I*) and LMSW223 (Δ*divIVA secA2 T123A*) during growth in BHI broth at 37°C. Membranes were stained with nile red. Scale bar is 2 µm. (C) Separation of secretome samples of the same set of strains by SDS-PAGE. The position of p60 (CwhA) is indicated. (D) Flagellar motility of the same set of strains on swarming agar after two days of incubation at 30°C. (E) Cell-to-cell spread of the same strains in 3T3 mouse fibroblast. (F) Full suppression of the Δ*divIVA* invasion defect by *secA2* mutations. The same set of strains as in panel A was used to infect HepG2 hepatocytes. Values are expressed relative to wild type. Average values and standard deviations are shown. The asterisk marks a statistically significant difference (*P*<0.01, *t*-test with Bonferroni-Holm correction, n. s. – not significant). (G) Partial suppression of the intracellular replication defect of the Δ*divIVA* mutant by *secA2* mutations. J774 mouse macrophages were infected with the same strains as above and intracellular bacteria were enumerated right after infection (0 h p. i.) and six hours later (6 h p. i.). Average values and standard deviations were calculated from experiments performed in triplicates. Statistically significant differences are indicated by asterisks (*P*<0.01, *t*-test with Bonferroni-Holm correction).

## Discussion

Genes that ensure separation of daughter cells following cell division are known determinants of *L. monocytogenes* virulence. Mutants lacking these genes or combination thereof form long chains of unseparated cells and show multiple virulence defects [17, 26–28, 31]. We here show that cell chaining generally does not explain these defects. First, depletion of EzrA, an early cell division protein acting as a membrane anchor for the Z-ring [35], did not impair invasion or intracellular growth, despite the formation of long filaments. EzrA-depleted cells even managed to spread from cell to cell, albeit with reduced efficiency, a capability, which is completely lost in the absence of *divIVA* [17]. The filaments of EzrA-depleted cells (approx. 13 μm long, see above) are comparable in length to the cell chains of the Δ*divIVA* mutant (approx. 15 cells per chain, at 1 to 1.5 μm in length). Thus, the length of the infecting particle alone cannot be the determining parameter. Second, introduction of a constitutively active *prfA\** allele was dominant over the Δ*divIVA* invasion defect: It not only restored wild-type invasion efficiency but even increased the invasion rate of the chain-forming Δ*divIVA prfA\** strain to the elevated level of the *prfA\** single mutant. The fact that cell chains can invade hepatocytes as efficiently as rods, provided that PrfA-dependent virulence genes are overexpressed, demonstrates that chain formation does not generally preclude invasion. Furthermore, it suggests that either the mechanisms regulating PrfA activity or the function of PrfA-dependent invasion factors may be impaired in the Δ*divIVA* mutant. The first possibility is unlikely, as the Δ*divIVA* mutant displays normal hemolysis, normal phospholipase activity and normal accumulation of ActA on its surface [17] suggesting wildtype-like virulence gene expression.

Remarkably, the Δ*divIVA* deletion was dominant over the *prfA*\* mutation in terms of intracellular replication in macrophages, indicating that this aspect of the Δ*divIVA* mutant’s virulence defect is independent of PrfA-dependent processes. In contrast, mutations in *secA2* partially restored this defect. The two suppressor mutations in *secA2* identified in this study affect the nucleotide binding domain of SecA2, restore autolysin secretion and consequently suppress cell chaining. One of them (*secA2 T123A*) is known to increase the ATPase activities of *B. subtilis* SecA in its basal and activated states [48], suggesting that DivIVA could act as an activator of SecA2’s ATPase activity. In the Δ*divIVA* mutant, the CwhA and NamA preproteins are synthesized but not recruited to septal SecA2, and are therefore not secreted [17]. This is in good agreement with the hypothesis of DivIVA as an activator of the SecA2 ATPase, particularly because ATPase activities of SecA proteins need to become activated for protein secretion, which is also achieved by preprotein binding or secretion pore assembly [49]. However, the intracellular replication defect and the inability of the Δ*divIVA* mutant to spread from cell to cell are only partially suppressed by either of the *secA2* mutations. This partial suppression may indicate that ATPase activation by these mutations is insufficient during infection of macrophages and fibroblasts, or that additional, SecA2-independent processes crucial for intracellular replication are impaired in the absence of DivIVA. The Δ*divIVA* mutant was found to be partially decorated with host actin intracellularly indicating that it escapes the phagosome and reaches the host cell cytoplasm, just as described for Δ*cwhA* and Δ*secA2* mutants [26, 50]. This finding aligns with observations that Δ*hly* and Δ*prfA* mutants, which are completely unable to escape the phagosome [51, 52], are more efficiently cleared by macrophages. In contrast, the Δ*divIVA* mutant survives in the host cell cytoplasm but replicates either not at all or only slowly, as suggested by the higher intracellular bacterial titers measured right after and six hours post-infection compared to Δ*hly* and Δ*prfA* mutants (Fig. 6B, Fig. 7B) [39]. The reason why the Δ*divIVA* mutant fails to replicate in the cytosol remains unknown. A delayed phagosomal escape could be an explanation and would be consistent with the reduced fraction of Δ*divIVA* cells decorated with actin. Additionally, the Δ*divIVA* mutant may be more susceptible to autophagosomal degradation [53], or it may exhibit auxotrophy for certain cytosolic nutrients, the uptake of which may be impaired due to the absence or malfunctioning of specific transporters. What seems more likely, however, is that the surface of the Δ*divIVA* mutant is altered in a way that affects the function of surface-anchored proteins including ActA (actin recruitment) and internalins (invasion). This would be consistent with the observation that restoration of autolysin secretion suppresses invasion and replication defects. To test these hypotheses, isolation and analysis of further Δ*divIVA* suppressors using experimental infections as selection tool would be required.

Besides its significance for understanding the role of DivIVA in virulence, our work also confirms the essentiality of EzrA and MreB, as demonstrated in previous studies [38, 39], and describes the phenotypes resulting from their depletion. The rod shape collapses when MreB, the main scaffold component of elongasomes responsible for peptidoglycan synthesis at the lateral cell wall, is depleted. This leads to the formation of coccoid cells with highly granular to particulate wall surface in larger zones, which are likely composed of folded peptidoglycan. In contrast, EzrA depletion prevents cytokinesis and leads to the formation of non-septated filaments. While depletion of both proteins does not interfere with invasion and intracellular replication, actin tail formation – and thus cell-to-cell spread – are affected. This demonstrates that morphological factors play a role in actin tail formation but further confirms that invasion and intracellular replication are independent of the rod shape.

## Materials and Methods

### Bacterial strains and growth conditions

All *L. monocytogenes* strains are listed in Table 1. Strains were cultivated in BHI broth or on BHI agar plates. Antibiotics and supplements were added at the following concentrations: erythromycin (5 µg/ml), kanamycin (50 µg/ml), X-Gal (100 µg/ml) or IPTG (as indicated). Growth in liquid medium was monitored either manually by measuring the optical density at 600 nm or automatically by recording the absorbance at 600 nm of bacterial cultures cultivated in microtiter plates using Tecan plate readers. *Escherichia coli* TOP10 was used as standard cloning host [54].

**Table 1:**
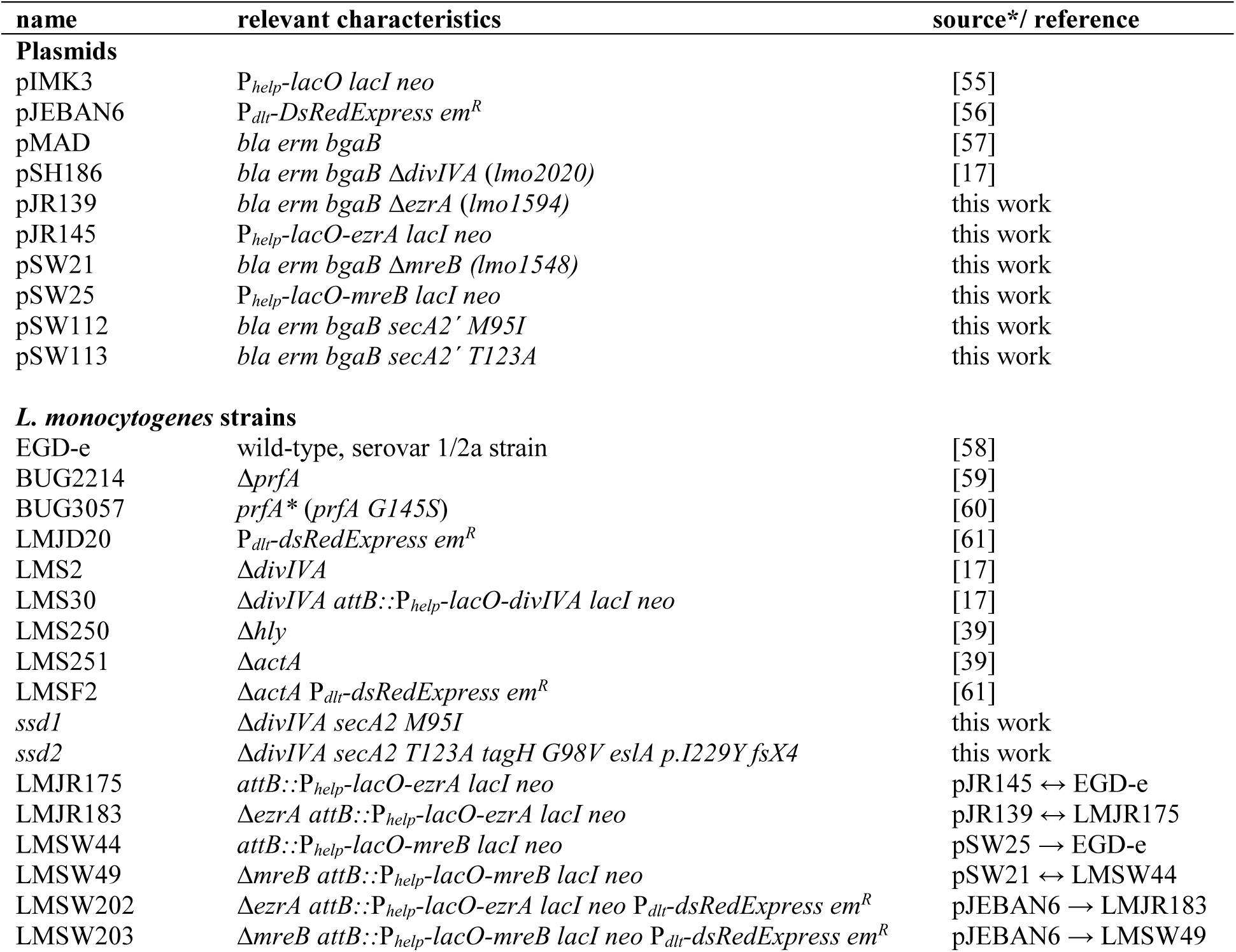
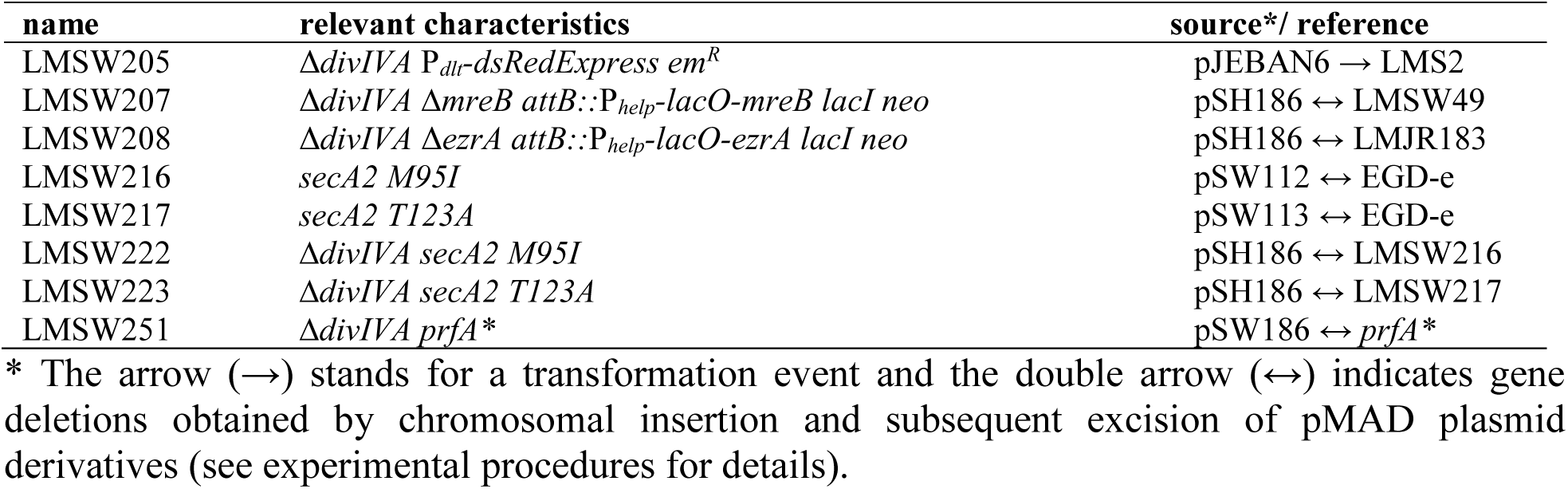
Plasmids and *L. monocytogenes* strains used in this study.

### General methods, manipulation of DNA and oligonucleotide primers

Standard methods were used for transformation of *E. coli* and isolation of plasmid DNA [54]. Transformation of *L. monocytogenes* was carried out as described by others [55]. Restriction and ligation of DNA was performed according to the manufactureŕs instructions. All oligonucleotides are listed in Table 2.

**Table 2:**
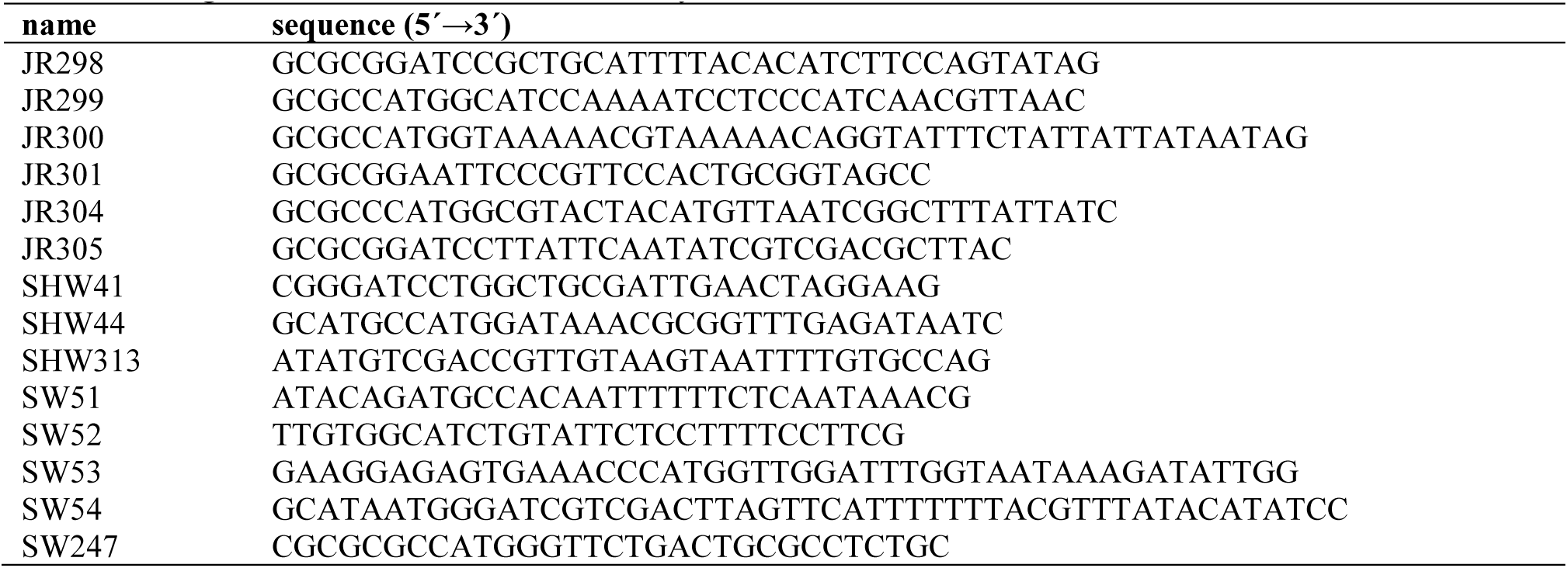
Oligonucleotides used in this study.

### Construction of plasmids and *L. monocytogenes* strains

All plasmids are listed in Table 1. Plasmid pJR139 was constructed for deletion of *ezrA*. To this end, *ezrA* up- and downstream fragments were amplified with JR298/JR299 and JR300/JR301, respectively, cut with NcoI and fused together by ligation. The desired Δ*ezrA* fragment was amplified from the ligation mixture using JR298/JR301 and cloned into pMAD using BamHI/EcoRI. Plasmid pJR145 was generated for IPTG-inducible expression of *ezrA*. It was obtained by amplification of *ezrA* using JR304/JR305 and cloning of the resulting fragment into pIMK3 after BamHI/NcoI digestion.

Plasmid pSW21 was constructed for deletion of *mreB*. For this purpose, *mreB* up- and downstream fragments were amplified using the oligonucleotides SHW41/SW52 and SHW44/SW51, respectively, and fused together by SOE-PCR. The resulting fragment was then cloned into pMAD using BamHI/NcoI. Plasmid pSW25 was constructed for IPTG-inducible *mreB* expression. To this end, the *mreB* gene was amplified from chromosomal DNA using SW53/SW54 and cloned into pIMK3 with NcoI/SalI.

For introduction of the M95I and T123A mutations into *secA2*, we constructed plasmids pSW112 and pSW113, respectively. To this end, N-terminal fragments of the *secA2* genes from strains *ssd1* and *ssd2* were amplified from chromosomal DNA using the oligonucleotides SW247/SHW313 and the resulting PCR fragments were cloned into pMAD using NcoI/SalI.

Derivatives of pIMK3 were introduced into *L. monocytogenes* strains by electroporation and clones were selected on BHI agar plates containing kanamycin. Plasmid insertion at the *attB* site of the tRNA^Arg^ locus was verified by PCR. Likewise, plasmid derivatives of pMAD were transformed into the respective *L. monocytogenes* recipient strains by electroporation and genes were deleted or mutation introduced as described elsewhere [57]. All gene deletions were confirmed by PCR.

### Whole genome sequencing

Chromosomal DNA was isolated by mechanical disruption of cell suspensions using glass beads in a TissueLyser II bead mill (Qiagen, Hilden, Germany) and quantified using a Qubit dsDNA BR Assay kit and a Qubit fluorometer (Invitrogen, Carlsbad, CA, USA). Libraries were prepared using the Nextera XT DNA Library Prep Kit and sequenced on a NextSeq sequencer in 2 x 150 bp paired end mode. Reads were mapped to the genome of *L. monocytogenes* strain EGD-e (NC_003210.1) [58] as the reference using Geneious (Biomatters Ltd.). Sequence variations were detected using the Geneious SNP finder tool. Genome sequences of strains sequenced in this study were deposited at the European Nucleotide Archive (ENA, https://www.ebi.ac.uk/) under project accession number PRJEB100780.

### Microscopy

Samples (0.4 µL) were taken from exponentially growing bacterial cultures and transferred onto microscope slides covered with a thin agarose film (1.5%). Samples were air-dried, covered with a cover lid and analysed by phase contrast or fluorescence microscopy. Chromosomes were stained using DAPI as described [24], while membranes were stained by addition of 1 µL of nile red solution (100 µg mL^-1^ in DMSO) to 100 µL of culture and shaking for 10 min at 37°C, before the cells were subjected to microscopy. Images were taken with a Nikon Eclipse Ti microscope coupled to a Nikon DS-MBWc CCD camera and processed using the NIS elements AR software package (Nikon) or ImageJ. Scanning electron microscopy were performed essentially as described earlier [62].

### Isolation of protein fractions, SDS PAGE and Western blotting

Cells were harvested by centrifugation (13,000 rpm, 1 min in a table-top centrifuge), washed with ZAP buffer (10 mM Tris-HCl pH7.5, 200 mM NaCl), resuspended in 1 ml ZAP buffer also containing 1 mM PMSF and disrupted by sonication. Cell debris was removed by centrifugation and the resulting supernatant was considered as total cellular protein extract. Aliquots of these samples were separated by standard SDS polyacrylamide gel electrophoresis. Gels were transferred onto positively charged polyvinylidene fluoride (PVDF) membranes using a semi-dry transfer unit. MreB and DivIVA were immune-stained using polyclonal rabbit antisera recognizing *B. subtilis* MreB [40] and *B. subtilis* DivIVA [63], respectively, as the primary antibodies and an anti-rabbit immunoglobulin G conjugated to horseradish peroxidase as the secondary one. The ECL chemiluminescence detection system (Thermo Scientific) was used for detection of the peroxidase conjugates on the PVDF membrane in a chemiluminescence imager (Vilber Lourmat).

For isolation of exoproteomes, bacterial strains were cultivated in 30 ml BHI broth at 37°C and grown to an OD_600_ of 1.0. Cells were pelleted by centrifugation and the secreted proteins were precipitated by the addition of trichloroacetic acid (10% wt/vol final concentration) to the supernatant fraction and over-night incubation at 4°C. Precipitated proteins were collected by centrifugation (6000 x g, 1 h, 4°C), and the dried protein pellet was resuspended in 8 M urea. Sample aliquots were separated by standard SDS polyacrylamide gel electrophoresis and stained with ROTI® Blue colloidal Coomassie stain (Carl ROTH, Karlsruhe, Germany).

### Hepatocyte invasion assays

Invasion of *L. monocytogenes* into HepG2 human hepatocytes (ATCC® HB-8065^TM^) was quantified as described earlier [64]. Briefly, 10^5^ hepatocytes were seeded into the wells of a 24 multi well plate and cultivated in DMEM + 10% fetal calf serum (FCS) overnight before they were infected with 2 x 10^5^ bacteria. Invasion was allowed to proceed during an incubation step at 37°C for 1 hour. Extracellular bacteria were first removed in a wash step with PBS and the remaining extracellular bacteria were killed by gentamicin addition. Eukaryotic cells were lysed right after infection using ice-cold PBS containing 0.1% Triton X-100, serial dilutions were plated on BHI agar plates and incubated overnight at 37°C for enumeration of bacterial cells.

### Macrophage infection

Macrophage infection experiments were performed as described [65]. Routinely, 3 x 10^5^ J774.A1 mouse ascites macrophages (ATCC® TIB-67^TM^) were cultivated in high glucose DMEM medium also containing 10% FCS for one day at 37°C in a 5% CO_2_ atmosphere. 5 x 10^4^ bacteria taken from overnight cultures were resuspended in 1 ml of DMEM not containing FCS and used as inoculum. Infection was allowed to proceed for 1 h. Afterwards, wells were washed with PBS and all remaining extracellular bacteria were killed by replacing the culture medium with fresh DMEM containing 40 µg/ml gentamicin and incubation for another hour. Thereafter, the wells were washed with PBS and then covered with fresh DMEM containing 10 µg/ml gentamicin and incubated at 37°C in a 5% CO_2_ atmosphere. Sampling was performed directly after infection and 6 h later by lysing the cells in 1 ml of ice-cold PBS containing 0.1% Triton X-100. For determination of bacterial cell counts, serial dilutions were plated on BHI agar plates.

### Microscopy of infected macrophages

3 x 10^5^ J774.A1 mouse ascites macrophages were seeded into the wells of a 24-well tissue culture plate containing coverslips and DMEM + 10% FCS as the culture medium and incubated in a 5% CO_2_ atmosphere at 37°C. Cells were infected as outlined above. Four hours after infection, cells were washed with PBS and then fixed with 4% paraformaldehyde for 20 min at room temperature. Coverslips were washed with PBS and then incubated in blocking/permeabilization buffer (2% BSA, 2% goat serum, 0.1% Triton X-100 in PBS) for 30 min at room temperature. The cover slip was dried and 20 µl of phalloidin staining solution (1 µl of a 66 µM alexa fluor^TM^ 488-phalloidin solution in DMSO was diluted in 200 ml PBS for this) were added and incubated for 1 h in the dark at room temperature. The coverslip was then washed with PBS and dried. Finally, a drop of ProLong Gold antifade reagent with DAPI (Invitrogen) was dropped onto a microscope slide and the coverslip was placed on top. Samples were dried overnight and examined using a Nikon Eclipse Ti fluorescent microscope.

### Plaque assays

Analysis of cell-to-cell spread was performed using 3T3-L1 mouse embryo fibroblasts (ATCC® CL-173T^M^) in plaque formation assays, as previously described [65]. Typically, 5 x 10^5^ fibroblast cells were seeded into the wells of a six well plate and cultured in DMEM supplemented with 10% newborn calf serum. After three days of incubation, cells were infected with an inoculum of 2, 4 or 10 μl each containing 2 x 10^6^/ml *L. monocytogenes* cells. Plaques were visualised three days post infection using neutral red staining.

### Hemolysis, phospholipolytic activity and motility assays

Hemolytic activity was assessed using the CAMP test [66]. For this purpose, *Staphylococcus aureus* SG511 was streaked perpendicularly to the *L. monocytogenes* strains to be tested on Mueller-Hinton agar plates containing 5% sheep blood. Zones of hemolysis were documented by photography after overnight incubation at 37 °C. An egg-yolk plate assay was employed to compare the phospholipolytic activity of *L. monocytogenes* strains [67]. For this purpose, one egg yolk (15 ml) was resuspended in 15 ml of sterile PBS buffer. The suspension was then mixed with 200 ml of molten LB agar containing 0.2% (w/v) activated charcoal and poured into Petri dishes.

*L. monocytogenes* strains were cultured on these plates, and zones of phospholipolysis became visible after two days of incubation at 37°C. To compare flagellar motility, *L. monocytogenes* strains were stab-inoculated on LB soft agar plates containing 0.3% agar. The plates were then incubated for 2 days at 30°C and swarming zones were photographed.

## Acknowledgements

We thank Birgitt Hahn for excellent technical assistance and Jeff Errington (University of Sydney) for kindly sharing the MreB antiserum.

## SUPPORTING INFORMATION CAPTIONS

**Figure S1:** Surface structure of *L. monocytogenes divIVA*, *ezrA* and *mreB* mutants.

Micrographs showing scanning electron microscopy images of *L. monocytogenes* strains EGD-e (wild type), LMS2 (Δ*divIVA*), LMJR183 (i*ezrA*) and LMSW49 (i*mreB*) from the same experiment as shown in Fig. 4 but at higher magnification. Strains were cultivated in BHI broth at 37°C to mid-logarithmic growth. For the IPTG-dependent strains, the preculture contained IPTG, but the inducer was omitted in the main culture.

**Figure S2:** Growth of IPTG-dependent *mreB* and *ezrA* mutants that also lack *divIVA*.

Growth of *L. monocytogenes* strains LMSW207 (Δ*divIVA* i*mreB*, A) and LMSW208 (Δ*divIVA* i*ezrA*, B) in BHI broth ± 1 mM IPTG. The inoculi for both growth experiments were taken from precultures cultivated in the presence of IPTG so that the presented experiments should be compared to the 1^st^ depletions shown in Figs. 1A and 2A. Average values and standard deviations from three independent repetitions are shown. Growth of all strains was measured in parallel but divided into two diagrams for better clarity.

**Figure S3:** Manifestation of EzrA and MreB depletion phenotypes during macrophage infection. J774 mouse macrophages were infected with *L. monocytogenes* strains LMJD20 (wt), LMSW202 (i*ezrA*), LMSW203 (i*mreB*) and LMSW205 (Δ*divIVA*) expressing DsRedExpress pre-grown with IPTG (where necessary) while the infection itself was carried out without IPTG for EzrA and MreB depletion. Pictures were taken four hours post infection. Phase contrast, blue channel (DAPI stained host cell nuclei), red channel (fluorescent bacteria) and merged images are shown.

**Figure S4:** Phenotype of the Δ*divIVA prfA\** mutant.

Hemolytic activity on sheep blood agar, phospholipolytic (PL) activity on egg-yolk agar, swarming on soft agar and cellular morphology after nile red staining (from left to right) was determined for *L. monocytogenes* strains EGD-e (wt), LMS2 (Δ*divIVA*), BUG2214 (Δ*prfA*), BUG3057 (*prfA\**) and LMSW251 (Δ*divIVA prfA\** ).

