## Supplementary figures S1-S4 for "On the role of cell chaining in the attenuation of a *Listeria monocytogenes divIVA* mutant"

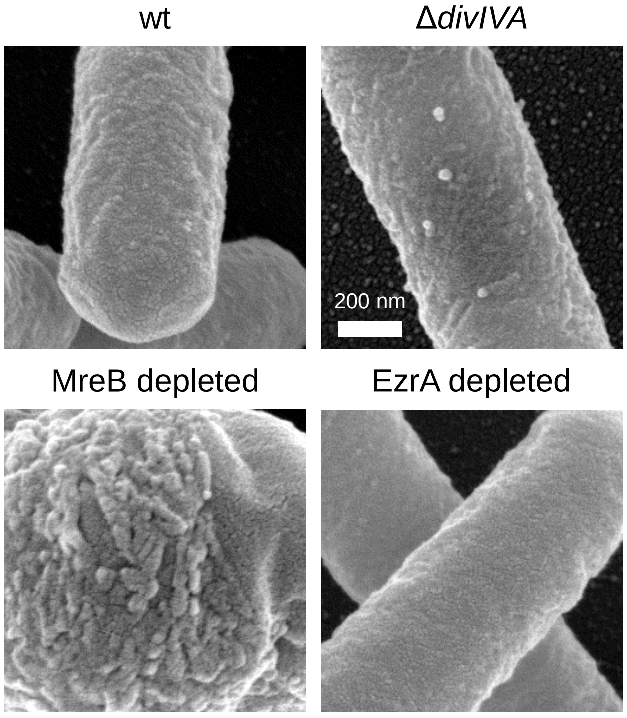


**Figure S1:** Surface structure of *L. monocytogenes divIVA*, *ezrA* and *mreB* mutants.
Micrographs showing scanning electron microscopy images of *L. monocytogenes* strains EGD-e (wild type), LMS2 (Δ*divIVA*), LMJR183 (i*ezrA*) and LMSW49 (i*mreB*) from the same experiment as shown in Fig. 4 but at higher magnification. Strains were cultivated in BHI broth at 37°C to mid-logarithmic growth. For the IPTG-dependent strains, the preculture contained IPTG, but the inducer was omitted in the main culture.

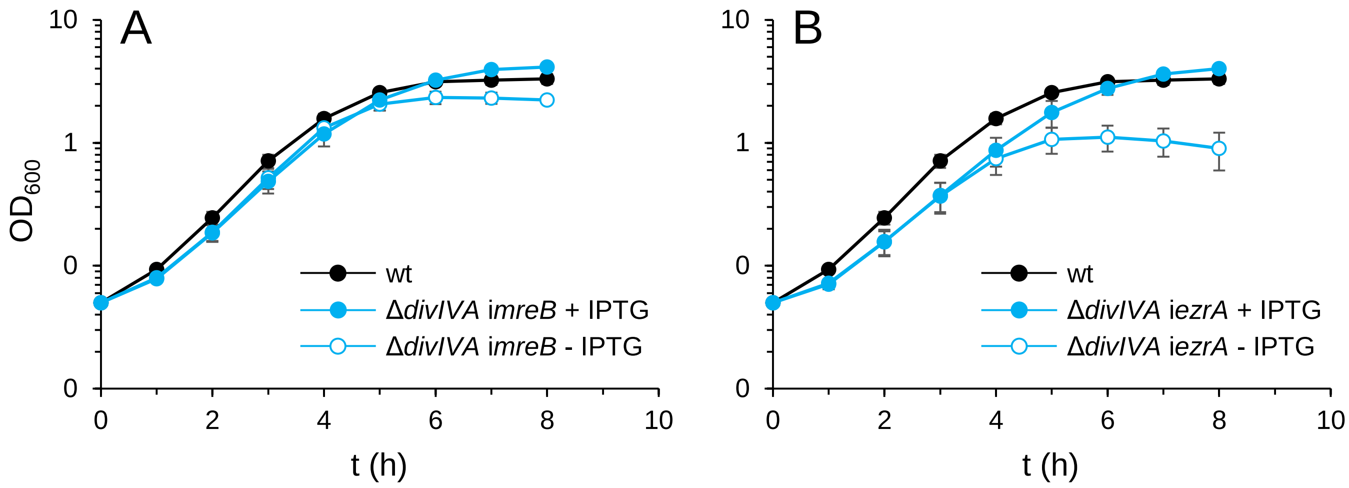


**Figure S2:** Growth of IPTG-dependent *mreB* and *ezrA* mutants that also lack *divIVA*.
Growth of *L. monocytogenes* strains LMSW207 (Δ*divIVA* i*mreB*, A) and LMSW208 (Δ*divIVA* i*ezrA*, B) in BHI broth ± 1 mM IPTG. The inoculi for both growth experiments were taken from precultures cultivated in the presence of IPTG so that the presented experiments should be compared to the 1st depletions shown in Figs. 1A and 2A. Average values and standard deviations from three independent repetitions are shown. Growth of all strains was measured in parallel but divided into two diagrams for better clarity.

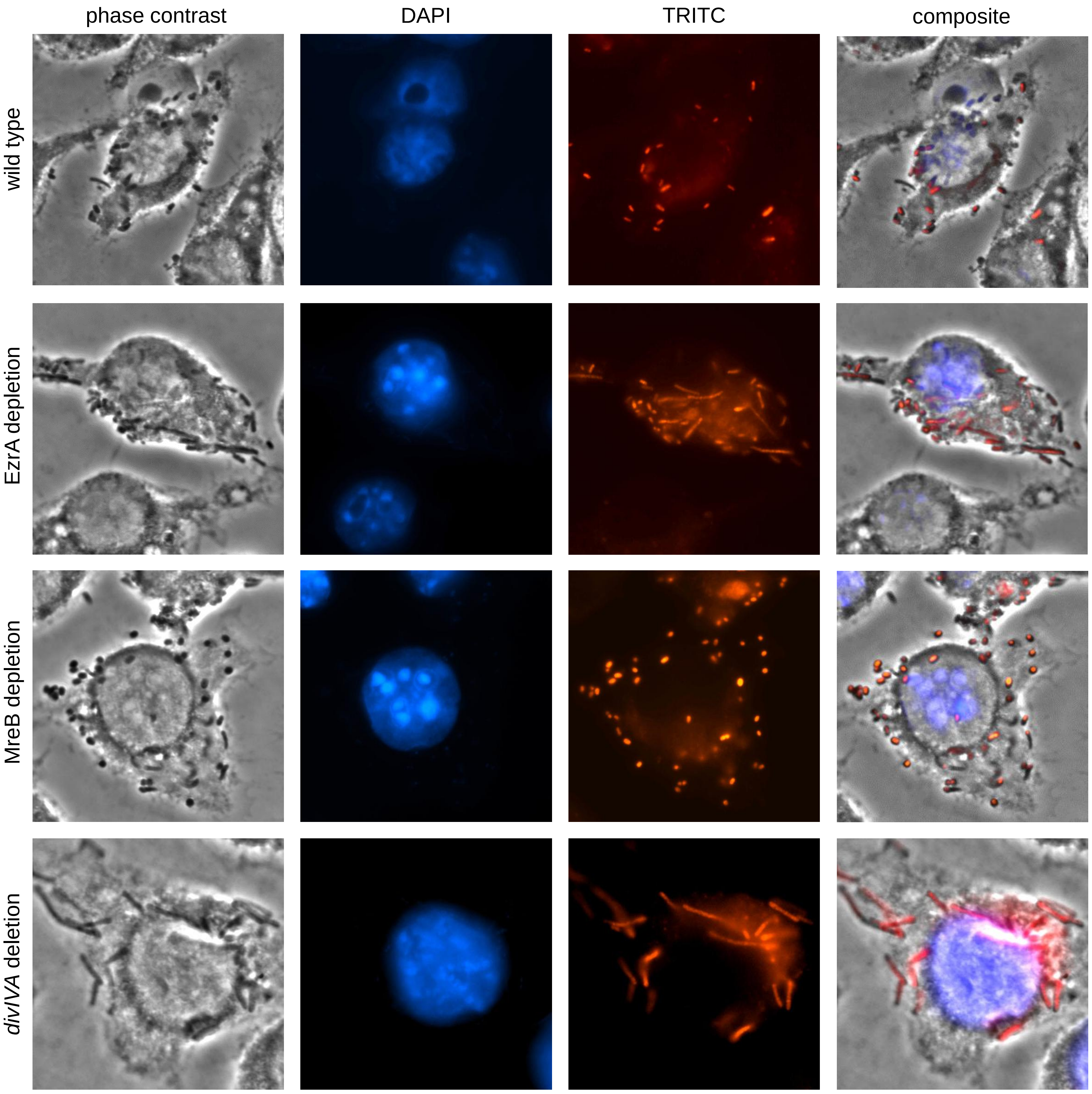


**Figure S3:** Manifestation of EzrA and MreB depletion phenotypes during macrophage infection.
J774 mouse macrophages were infected with *L. monocytogenes* strains LMJD20 (wt), LMSW202 (i*ezrA*), LMSW203 (i*mreB*) and LMSW205 (Δ*divIVA*) expressing DsRedExpress pre-grown with IPTG (where necessary) while the infection itself was carried out without IPTG for EzrA and MreB depletion. Pictures were taken four hours post infection. Phase contrast, blue channel (DAPI stained host cell nuclei), red channel (fluorescent bacteria) and merged images are shown.

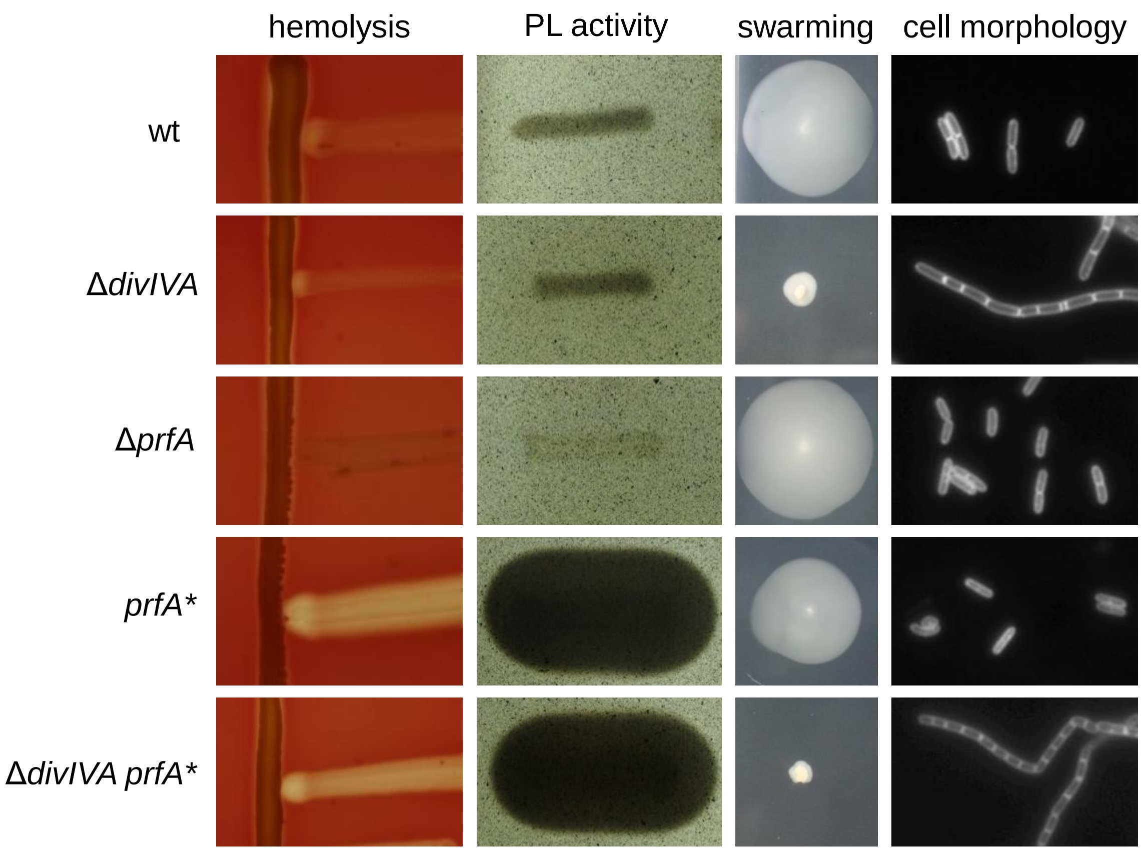


**Figure S4:** Phenotype of the Δ*divIVA prfA** mutant.
Hemolytic activity on sheep blood agar, phospholipolytic (PL) activity on egg-yolk agar, swarming on soft agar and cellular morphology after nile red staining (from left to right) was determined for *L. monocytogenes* strains EGD-e (wt), LMS2 (Δ*divIVA*), BUG2214 (Δ*prfA*), BUG3057 (*prfA**) and LMSW251 (Δ*divIVA prfA** ).
